# Splendid isolation with migration: Diversity dynamics of South American mammals

**DOI:** 10.1101/2025.11.28.691169

**Authors:** Kateryn Pino, Enrique Rodríguez-Serrano, Juan D. Carrillo, Rebecca B. Cooper, Daniele Silvestro

## Abstract

Biodiversity dynamics following encounters between long-isolated faunas provide natural experiments to examine how diversity dependence and ecological interactions shape diversification patterns at macroevolutionary time scales. The Cenozoic arrival of African and North American mammals in South America likely had transformative effects on one of the world’s most unique and previously isolated continental faunas. However, the extent and persistence of these effects remain debated. Here, we assess the impact of immigrant mammal groups on native South American mammal diversity by combining an extensive fossil dataset (6,214 occurrences; 1,739 species) with deep learning methods and time-series analyses to model dynamics of mammalian biodiversity throughout the Cenozoic. The arrival of African and North American lineages increased the continent’s overall species richness but also contributed to a decline in native diversity. Additionally, our models suggest that the negative effect of immigrant lineages on native diversity was strongest immediately after their arrival and initial diversification, but this effect lessened over time. Overall, these findings support a scenario where immigration simultaneously enriched South American faunas and triggered a time-decaying replacement of native lineages, reshaping continental biodiversity.

## Introduction

Throughout the history of life, major clades have waxed and waned; once-dominant groups have disappeared, while others have risen to take their place^1^. These patterns of clade turnover, observed at both global and regional scales, have shaped ecosystems throughout the Phanerozoic^2,3^. Such clade turnover is ultimately driven by macroevolutionary processes like speciation, extinction, and, at regional scales, dispersal. These dynamics are influenced by both physical factors and biotic interactions, including competition and predation^4^. Among these, competition has been proposed as a key factor driving clade replacement, particularly through diversity-dependent processes^5–11^. Diversity-dependence refers to the impact of existing diversity on macroevolutionary dynamics, whereby high levels of diversity can constrain speciation in ecologically saturated environments^12,13^ but may also elevate extinction risk via competitive exclusion among ecologically similar clades^8,14^.

Among the processes contributing to biotic replacement, those involving the arrival of lineages that initially diversify and evolve in isolation and subsequently coexist due to dispersal or changes in geographic connectivity are particularly revealing^1,15^. These biogeographic events provide unique opportunities to investigate how competition and ecological interactions influence diversification patterns at macroevolutionary scales^15,16^. The arrival of immigrant lineages into a region may alter the diversification dynamics of both immigrants and natives, either by expanding overall diversity or by inducing turnover through diversity-dependent interactions^8,10,17^. If the region supports more species, immigrants can enrich its diversity as they find ecological opportunities to diversify^18,16^. Conversely, the arrival of new lineages to the region could be counterbalanced by the extinctions of native organisms, as expected by diversity-dependent processes^13,20^. Furthermore, these biogeographic phenomena enable us to investigate whether negative interactions and/or diversity-dependent processes persist or weaken over time. Over the evolutionary timescale, the strength of interspecific competition can decrease through spatial segregation, character displacement, and niche specialization^21–23^. Therefore, the intensity of negative biotic interactions between native and immigrant taxa can diminish over time as ecological interactions restructure, with descendants of the new lineages evolving on the new continent and native lineages adapting, ultimately reaching a new equilibrium^13,16^.

The evolution of South American (SA) mammals over the Cenozoic provides a compelling natural experiment for studying biotic replacement patterns and the processes involved^7,11,18,24,25^. Mammalian faunas in this continent evolved through extended periods of biogeographic isolation punctuated by rare dispersal events during the Paleogene, followed by increased connectivity during the Great American Biotic Interchange (GABI) with the formation of the Panama Isthmus^7,11,18,26,27^. Simpson proposed a *three-stratum* model to describe the pattern of episodic replacement of the native SA mammal faunas by the arrival of new lineages from Africa and North America^18,28^. The first *stratum* comprises the oldest native SA mammal faunas, including three main groups: metatherians, xenarthrans, and SA native ungulates (SANUs). The second *stratum* includes long-distance dispersal lineages from Africa, i.e., primates and caviomorph rodents during the Late Eocene^26–31^ or Early Oligocene^32^. Finally, the third *stratum* involves the dispersal of mammalian faunas from North America during the GABI from the Late Miocene to the Pleistocene. This event includes the appearance of placental carnivores, perissodactyls, artiodactyls, proboscideans, cricetid rodents, lagomorphs, soricids, and sciurids in the SA fossil record for the first time.

These faunal shifts have been interpreted through the lens of diversity-dependence dynamics and have inspired extensions of the island biogeography theory to continental settings^7,11,20^. The arrival of ecologically similar taxa likely triggered competition and/or predation, potentially driving the extinction of native forms, such as predator-naïve prey confronted by placental carnivores, leading to immigration-induced turnover^11,24,33,34^. Replacement by competition is expected, particularly among ecological equivalents—taxa that occupy similar adaptive zones and trophic roles^8,12,35^, such as the turnover in the carnivore and herbivore guilds of SA mammalian faunas^11,18,28,34,36^. Alternatively, a passive displacement might also explain this pattern of faunal turnover^37,38^, with immigrants filling the ecological vacancies left by natives diminished by physical factors. Environmental changes (e.g., habitat change, Andes uplift, and climate shifts) have been suggested as the main driver behind clade replacement in SA mammalian faunas, with immigrant and native lineages responding differently to these changes^24,36–42^.

Here, we investigate how the arrival of African and North American lineages into South America influenced native diversity and whether these events led to faunal enrichment and/or replacement. To this aim, we first conceptualize the expected patterns of diversity trajectories under three evolutionary hypotheses. First, under an ecological opportunity hypothesis, we postulate that the arrival of immigrants enriches the SA fauna as new lineages find ecological spaces previously unoccupied by the natives to diversify. This leads to an overall increase in diversity, while the native diversity remains unaffected (Fig. 1a). Second, under a competitive replacement hypothesis, the arrival of immigrants drives turnover and replacement through negative biotic interactions affecting the local fauna. In this case, the overall diversity remains unchanged as native diversity decreases while the diversity of immigrants increases (Fig. 1b). Third, as hypotheses 1 and 2 are non-mutually exclusive, a combination of them is plausible. Under this scenario, the arrival of immigrant lineages enriches the recipient fauna, but the native diversity is affected and partially replaced by the immigrants (Fig. 1c). We test these hypotheses in a quantitative framework, analyzing the Cenozoic fossil record of SA mammals using statistical and deep learning methods.

**Figure 1.**
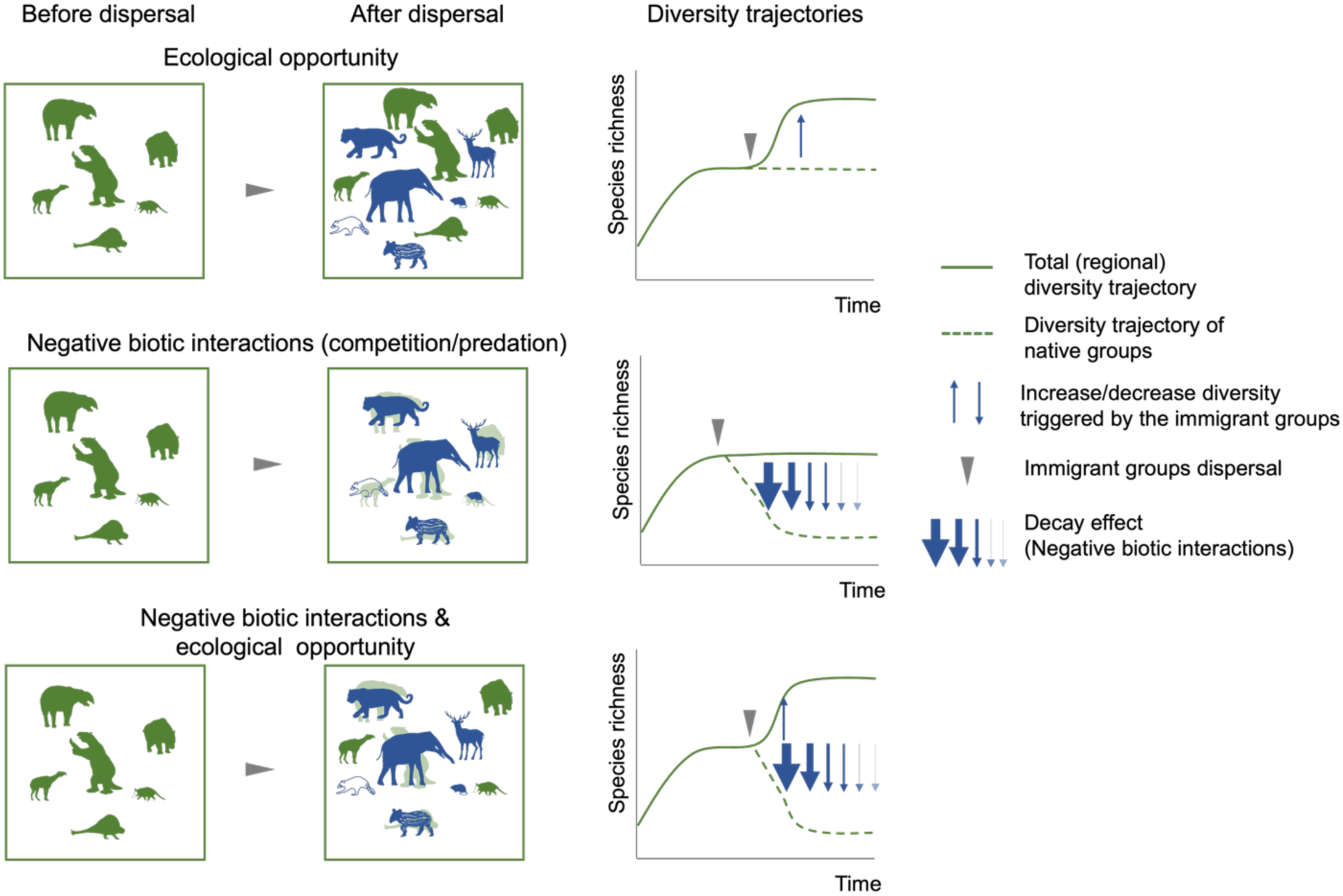
Evolutionary hypotheses of biotic dispersal and replacement during encounters between biotas with long-separated evolutionary histories. If immigrants encounter an unsaturated recipient fauna or are able to utilize an open ecological space, they will find an opportunity to diversify, resulting in the enrichment or increase in overall diversity without affecting the pre-existing fauna. Conversely, if the recipient fauna is saturated and ecological space or adaptive zones are occupied by native taxa, then immigrants drive turnover through negative biotic interactions, and the overall diversity remains unchanged, as immigrants replace native diversity. Finally, the immigrants enrich the recipient fauna, but partial duplication by native and immigrant taxa in the ecological space is inevitable.

## Results

### Diversity patterns of South American mammals throughout the Cenozoic

Through the stochastic simulations and deep learning models implemented in DeepDive^43^, we estimated the diversity trajectories of SA mammals over the Cenozoic, while accounting for temporal, spatial, and taxonomic biases in the fossil record. We found that the diversification of immigrant groups, primates, and caviomorphs from Africa during the Eocene-Oligocene and multiple North American lineages from the Late Miocene to the Pleistocene consistently led to overall net increases in mammalian species richness in the continent, reaching its zenith during the Early Pleistocene (mean 1282 species, 65% CI = [1058, 1506]). Yet, these migration events also coincided with the decline of the diversity of SA native lineages.

South American native diversity increased steadily, with some fluctuations, during the Paleocene and Early Eocene, reaching around 754 mean species, 65% CI = [636, 877] by the Middle Eocene (40 Ma, Bartonian). Diversity then gradually declined by about 36% until the end of the Late Oligocene (∼24 Ma). The initial phase of this decline coincided with the arrival and subsequent diversification of primates and caviomorph rodents during the Late Eocene. Throughout the Neogene, SA native species diversity underwent significant fluctuations. It rebounded in the Early Miocene, followed by a steep decline before undergoing another recovery in the Middle Miocene. However, diversity began to decline rapidly around 7 million years ago and has not recovered since, losing 73% of its diversity until reaching its current state. Interestingly, this sharp decline overlaps temporally with the diversification of multiple North American lineages that arrived in SA since the end of the Serravallian (Fig. 2).

**Figure 2.**
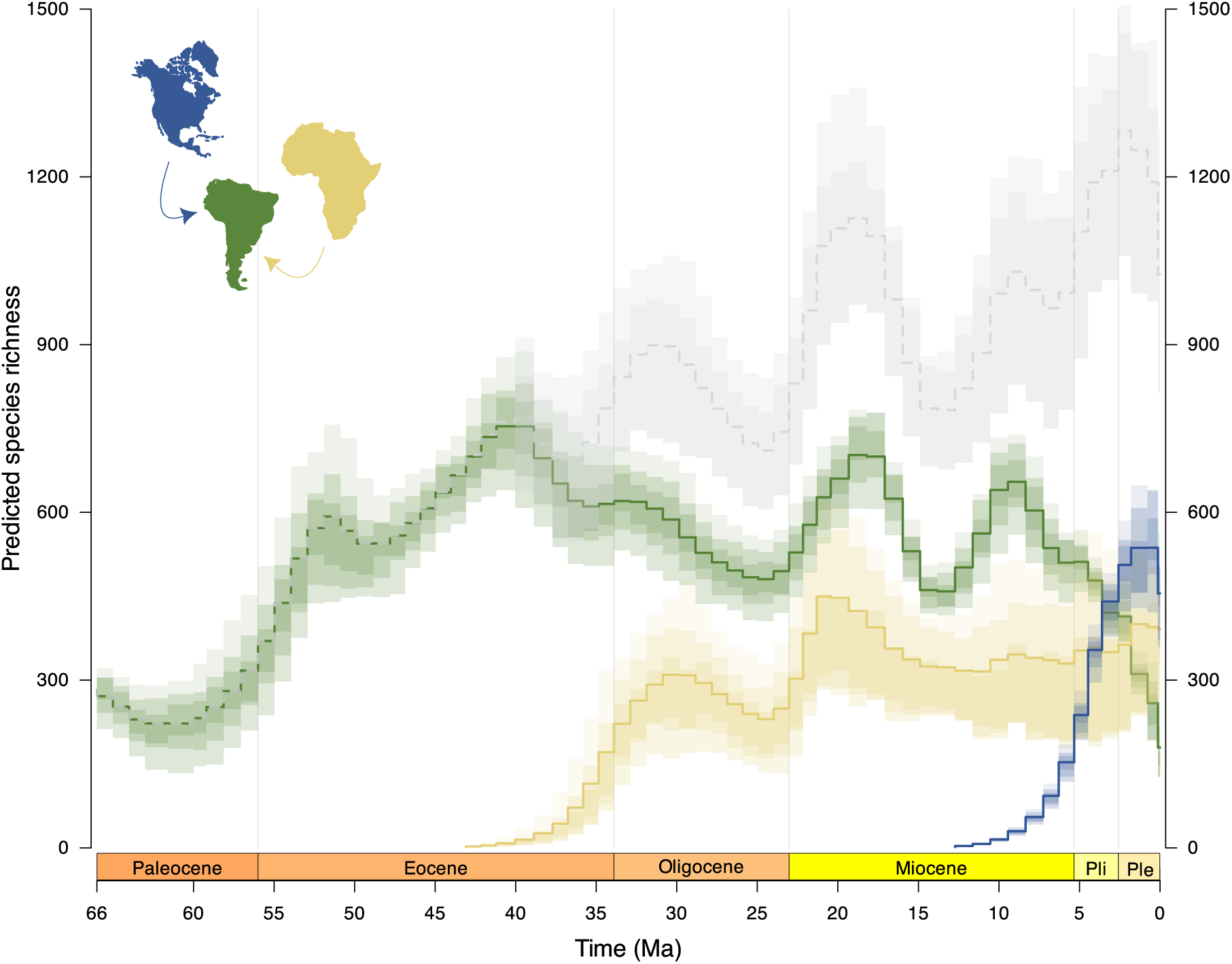
Diversity trajectories of the combined mammal groups per origin. Native groups from South America (green) (metatherians, Xenarthrans, and SANUs), newly arrived groups from Africa (yellow) (Caviomorpha and Primates), and North America (blue) (Carnivora, Cricetidae, Artiodactyla, Perissodactyla, and Proboscidea). The diversity trajectory of all mammals in South America is shown in gray. Curves display the mean 50%, 70%, and 65% confidence intervals.

### Diversity trajectories of the major groups of South American mammals

Non-sparassodont metatherian diversity (i.e., the combined species richness of Didelphimorphia, Paucituberculata, Microbiotheria, “Polydolopimorphia,” and “Ameridelphimorphians”; hereafter referred to as metatherians) gradually increased during the Paleocene, reaching its highest species richness ∼51 Ma (mean of 317 species, 65% CI = [183-466]) during the Ypresian. Afterward, diversity declined sharply and continued to decrease until the beginning of the Priabonian (Late Eocene), when it began to accumulate again, maintaining some fluctuations through the remainder of the Neogene. However, the diversity level achieved in the Ypresian has never been recovered again (mean loss of 64% of species). The species richness of metatherian carnivores of the Sparassodonta group exhibits a different pattern. Diversity built up slowly during the Paleocene and most of the Eocene, reaching its peak species richness during the Bartonian (mean 44 species, 65% CI = [32, 57]) and Early Miocene (mean 58 species, 65% CI = [42, 72]). Species richness declined in the Priabonian (mean 47% species loss) but rebounded during the Oligocene; however, it began to decrease again, first slowly in the Middle Miocene and then crashing toward the end of the Miocene until it became extinct in the Plio-Pleistocene (Fig. 3).

**Figure 3.**
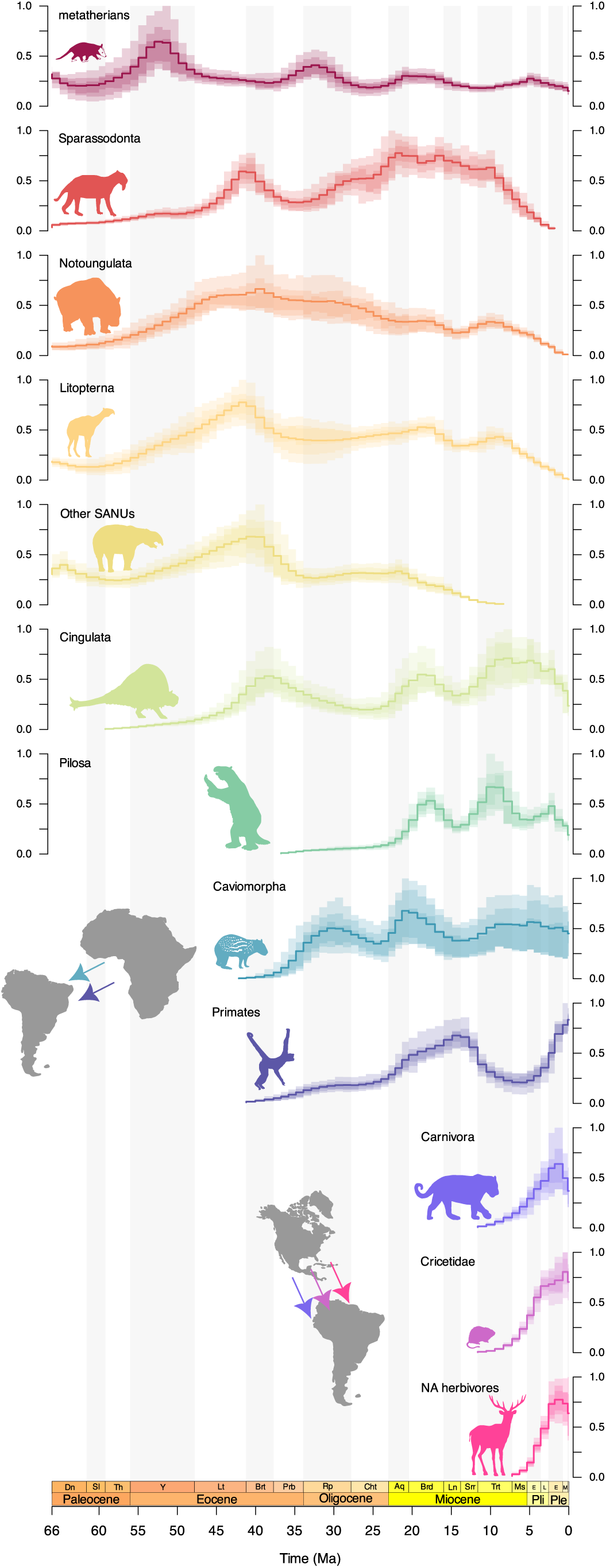
Diversity trajectories of South American mammal groups throughout the Cenozoic. Metatherians comprise Didelphimorphia, Paucituberculata, Microbiotheria, “Polydolopimorphia”, and the basal “Ameridelphimorphians” (*sensu* Goin 2022^44^). Other SANUs consist of Astrapotheria, Pyrotheria, Xenungulata, and Condylarthra, while North American (NA) herbivores include the orders Artiodactyla, Perissodactyla, and Proboscidea. Species richness values have been rescaled so that the maximum value in each 65% confidence interval equals 1. Representative silhouettes of the groups are sourced from phylopic.org.

The diversity trajectory of SANUs, including Notoungulata, Litopterna, and “Other SANUs” (i.e., the combined species richness of Astrapotheria, Pyrotheria, Xenungulata, and Condylarthra), exhibits similar diversity patterns, reaching their highest species richness in the Bartonian. Notoungulata species richness then steadily declined until the Serravallian, followed by a brief recovery phase during the Tortonian. In contrast, Litopterna and the “Other SANUs” experienced a rapid decline in the Late Eocene and remained stable until the Early Miocene, when the diversity of “Other SANUs” continued to decrease until its extinction in the Tortonian, whereas Litopterna and Notoungulata diversity began to diminish toward the end of the Miocene, ultimately going extinct in the Late Pleistocene (Fig. 3).

Cingulata diversity built up from the end of the Paleocene toward the end of the Bartonian, after which its species richness waned until the end of the Oligocene. Throughout the Early and Middle Miocene, Cingulata and Pilosa exhibit similar diversity trajectories, with species richness waxing in the Early Miocene and then waning afterward in the Middle Miocene before rebounding during the Tortonian when they reach their peak (mean 181, 65% CI = [103, 255] and 146 species, 65% CI = [83, 218]; respectively). Then, the species richness of Pilosa exhibits a decline phase followed by a brief recovery during the Late Miocene, while Cingulata diversity remains constant. Finally, both groups experience a rapid decline that begins in the Early Pleistocene and becomes more pronounced toward the Late Pleistocene (Fig. 3).

Diversity of immigrant groups exhibits a rapid rise following dispersal from Africa and North America (but see primates, Fig. 3). Caviomorph rodent species richness rose from the Late Eocene to the Early Oligocene and in the Early Miocene, while it diminished during the Late Oligocene and Middle Miocene, rebounding toward the Late Miocene and remaining constant since then. In contrast, primate diversity increased slowly until the Languian (Middle Miocene) before beginning to decline throughout the rest of the Miocene, to then rebound during the Pliocene and Pleistocene. North American lineages reached their zenith during the Early-Middle Pleistocene, after which the diversity of Carnivora and NA herbivores (i.e., the combined species richness of Artiodactyla, Perissodactyla, and Proboscidea) dropped slightly in the Late Pleistocene, while cricetids continued to increase (Fig. 3).

### Native and Immigrants’ time series correlation

To quantitatively assess our negative biotic interactions hypotheses (i.e., competition and predation) between native and immigrant groups, we use the diversity time series estimated with DeepDive to model multiple regressions. We also included climatic and environmental factors in our regressions to account for abiotic effects and use Akaike Information Criterion (AICc) to select the significant predictors. Our models estimated the extent to which the diversity trajectory of native mammals can be explained by changes in immigrant diversity and additional abiotic factors. We also included in the estimation a decay parameter describing biotic interactions that decline over time. Thus, when the estimated decay parameter equals zero, the interaction between immigrant and native lineages (as quantified by the regression coefficient) persists over time, while a decay parameter greater than zero indicates that the interaction fades over time (see Methods).

For the total diversity model, we found that the total diversity of immigrants from North America (i.e., combined species richness of NA herbivores, placental carnivores, and cricetid rodents) is negatively correlated with the total SA native diversity (i.e., combined species richness of Metatheria, Xenarthra, and SANUs) (model-averaged regression coefficient = -173.637; 65% CI = [-271.466, -75.808]) (Fig. 4; Table S4). Yet, we did not find an association with the total diversity from Africa (i.e., combined species richness of platyrrhines and caviomorphs). Although the top-ranking models (i.e., cumulative AICc weights > 0.65) include this predictor to explain the diversity dynamics of the total SA native diversity, the 65% confidence interval of its average coefficient overlapped zero (model-averaged regression coefficient -86.575; CI = [-182.634, 6.483]) (Fig. 4; Tables S3, S4). In addition, we found a pervasive positive association between the Andes and the SA native diversity in all our models. However, we did not find support for any association with the changes in the southern SA landscape, from dense vegetation to more open habitats (Fig. 4; Tables S4, S6, S8).

**Figure 4.**
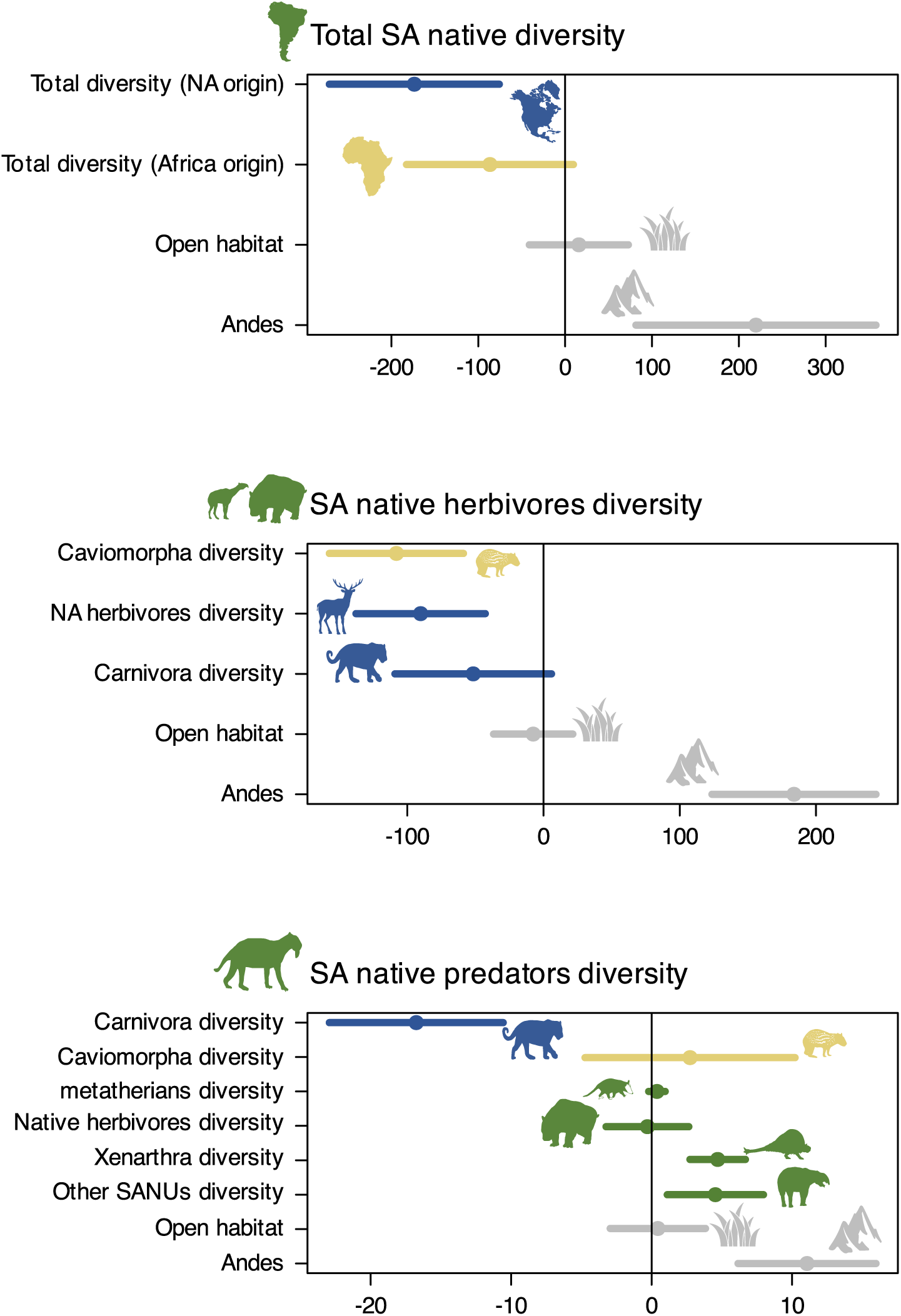
Model-averaged regression coefficients and confidence intervals for each predictor in our model hypothesis (see Methods). Arranged from top to bottom: total diversity-dependence, herbivore and predator-naïve prey, and predator competition hypotheses. Confidence intervals not including zero provide strong evidence of affecting the response variable, whereas those that include zero are considered uninformative (high uncertainty). Colors indicate the continent of origin for the groups: South America (green), North America (blue), and Africa (yellow). Representative silhouettes are sourced from phylopic.org.

Within the herbivore and predator guilds, we found that the diversity of immigrant groups is correlated with a decrease in native diversity, supporting the hypotheses of replacement by competition. Our averaged models indicate that the diversity of immigrant herbivores from Africa (i.e., species richness of Caviomorpha) (model-averaged regression coefficient = -60.166; 65% CI = [-137.642, -42.75]) and North America (i.e., combined species richness of Artiodactyla, Perissodactyla, and Proboscidea) (model-averaged regression coefficient = 107.68; 65% CI = [-157.186, -58.776]) had a negative impact on the diversity of SA native herbivores (i.e., combined species richness of Notoungulata and Litopterna). Moreover, the diversity of immigrant placental predators from North America (i.e., Carnivora species richness) is negatively correlated with SA native predators (i.e., Sparassodonta) (model-averaged regression coefficient = 107.68; 65% CI = [-157.186, -58.776]) (Fig. 4; Tables S7, S8) and positively associated with the diversity of their potential native prey, xenarthrans and “other SANUs” (model-averaged regression coefficient = 4.536; CI = [1.115, 7.664]; model-averaged regression coefficient = 4.704; CI = [2.73, 6.677], respectively) (Fig. 4; Table S8). The impact of immigrant diversity on native diversity is highest at the onset of its diversification and decreases over time. All the best-fitting models in our tests included a non-zero *decay effect* parameter, with values ranging from 0.016 to 0.048 (Tables S3, S5, S7; Figs. S2-S7). Thus, the impact of immigrant lineages on native clades was estimated to be strongest in the aftermath of the migration event, while declining over time (Figs. S2-S7).

We did not find support for the predator-naïve prey hypothesis (i.e., the arrival of placental carnivorans from North America explaining the diverse dynamics of native SA herbivore lineages), as the 65% confidence interval of the parameter estimates in the model averaging overlapped zero (model-averaged regression coefficient = -51.564; CI = [-106.044, 5.857]) (Fig. 4, Table S6).

## Discussions

Our results show that the arrival and subsequent diversification of mammalian lineages from Africa and North America into South America had profound effects on the evolutionary dynamics of mammal communities in the continent, while leading to sustained increases in overall species richness. Immigration events were also consistently accompanied by significant and long-lasting declines in the diversity of native South American clades, especially among herbivorous and carnivorous groups. Statistical modeling of diversity trajectories and their correlations supports the hypothesis that competitive interactions could influence these patterns of biotic replacement; however, the negative impact of immigrants decreases over time.

These findings reveal that biotic replacement in South America during the Cenozoic was a dynamic process likely influenced by diversity-dependent interactions between immigrant and native lineages. While the continent experienced substantial faunal enrichment following dispersal events, particularly during GABI, this overall increase in total richness masked a turnover of native taxa. The decline of native groups alongside the rise of ecologically similar immigrants, especially among herbivores and carnivores, highlights the role of competitive exclusion at macroevolutionary scales^4,45^. These results provide quantitative support for long-standing theoretical frameworks, such as Simpson’s *three-stratum* model and Webb’s and Marshall’s adaptation of island biogeography to continental contexts^7,11^, while also offering a revised temporal resolution to the dynamics of faunal replacement. Crucially, we show that the negative effects of immigrant clades on native species are strongest at the beginning of their diversification and weaken over time. Thus, SA clades appear to have been pushed to a lower level of diversity by immigrant lineages but later reached a new equilibrium that allowed them to coexist.

### Immigration led to the enrichment of continental diversity in South America during the Cenozoic

The immigration of taxa into previously isolated regions can trigger their radiation as they reach evolutionarily accessible resources or adaptive zones that may be underutilized by native taxa^12,14,46^, leading to the enrichment of biotas on continents and oceans^15,17,47,48^. Newly introduced lineages provide new opportunities for subsequent evolution once the dispersal pathways are disconnected or unavailable. Indeed, the isolation of geographic areas enhances speciation through vicariance and promotes an increase in biodiversity^17^. Furthermore, the success of immigrant lineages in a new region may result from the release from natural enemies (i.e., predators, competitors, disease agents, and parasites). This ecological release enables them to achieve greater diversity than their relatives in their region of origin^16^. For instance, the sigmodontine and caviomorph rodents experienced rapid radiation after their arrival on the SA continent, attaining considerably higher diversity than their sister groups in their origin regions (i.e., North America and Africa, respectively)^46,50^. Yet, the radiation of immigrant lineages in the new region could impact the native fauna, inducing turnover and replacement^16,17,24^, as also seen in present-day biological invasions. Invasion biology studies on birds and plants in oceanic islands at ecological time scales show endemic species often going extinct due to the arrival of immigrant lineages; yet, local diversity remains either unchanged or rises^51^. Moreover, a long-term study of tree communities at a continental scale has shown that the species richness of non-native species has increased, while that of native species has decreased^52^. Yet, native biomass and trait-and phylogenetic-diversity-independent measures have tended to remain stable. Furthermore, displaced native species are functionally and phylogenetically similar to non-native species; in contrast, native survivors tend to be functionally distinct from non-native invaders, suggesting an essential role for competitive exclusion and resource partitioning in community dynamics during biological invasions^52^.

### Immigration induced faunal turnover in South America

The episodic succession of SA mammal faunas follows a double-wedged diversity trajectory pattern, with native species decreasing while immigrant species increase simultaneously. This pattern, where one clade rises at the same time another falls, has been discussed as a possible outcome of ecological interactions between taxa from different clades that are ecologically similar^22,53,54^. The turnover and partial replacement of SA native clades by immigrant clades from Africa and North America can be attributed in part to competitive exclusion among ecological groups occupying the same trophic level within an adaptive zone, specifically among herbivores and carnivores (Fig. 4; Tables S6, S8).

The herbivore-adaptive zone of SA mammalian fauna was mainly dominated by SANUs, highly diverse ungulate-like herbivores that were almost exclusively found on the continent and were abundant during most of the Cenozoic, but are now entirely extinct^34,55,56^. The dominance of SANUs in the herbivore adaptive zone in South America was likely affected by the diversification of caviomorph rodents during the Late Eocene–Oligocene and then by North American herbivores (i.e., artiodactyls, perissodactyls, and proboscideans) during GABI in the Pliocene-Pleistocene. The most species-rich, abundant, and long-lived SANUs were notoungulates and litopterns, both of which survived until the Late Pleistocene-middle Holocene^56–58^. Many litopterns are broadly similar to modern ungulates in terms of limb morphology and, to a large extent, craniodental morphology^56,56^. For instance, locomotor adaptations of early litopterns and proterotheriids closely resemble those of artiodactyls and caviid rodents^60,61^. Moreover, typotherians, a diverse suborder of primarily small to medium-sized notoungulates (1-10 kg, although some mesotheriids reached ∼75 kg)^56,62^, have many craniodental similarities with rodents and rabbits (see ^56,63,64^ and reference therein). For instance, pachyrukhines exhibit a mosaic of traits, combining plesiomorphic characteristics found in ungulates with more derived traits that are convergent with rodents^65^. Therefore, the strong correlation between the rise in immigrant taxa and the decline in native taxa, along with the close morphological and ecological similarities between them, suggests that an active replacement between these herbivore clades may have occurred.

The carnivore guild of SA mammalian fauna was dominated by sparassodonts, an endemic and highly specialized clade of metatherians that occupied this guild from the Paleocene to the Late Pliocene^66,67^. Sparassodonts were replaced by placental carnivores that arrived from North America at the beginning of the GABI^18,28,34^. However, they differed in their morphology and ecology; placental carnivores exhibited higher disparity and diversity than sparassodonts^68–70^. The earliest fossil record for Carnivora lineages in South America is recorded in three localities in the Pampean region of Argentina, with inferred ages of ∼7 Ma (range 7.31-6.32 Ma)^71,72^. However, their arrival time on the continent has been estimated to be around 6.6 Ma (65% credible interval: 8.1–12.2 Ma) based on fossil occurrences and Bayesian analyses that consider the fossil sampling and preservation bias^36,73^. Thus, placental predators and metatherians could have overlapped temporally for ∼7 Myr (from the estimated arrival of Carnivora at ∼6.6 Ma to the extinction of Sparassodonta at ∼2.6 Ma). Our estimated diversity trajectories are in agreement with this time overlap; moreover, we found a significant correlation between the decline of sparassodont species richness and the rise in placental carnivore diversity during this time (Figs. 3, 4; Tables S7, S8).

Our results appear to contradict previous studies that rule out the hypothesis of replacement by competition between these predator clades^36,41,74^. They found that the decline of sparassodonts begins before the arrival of placental carnivores (∼17 Ma), when the extinction rate increases, exceeding the origination rate, resulting in a negative diversification rate since then. Our estimated diversity trajectories confirm a gradual decrease in sparassodont species richness that began at this time and continued until ∼6 Ma. Yet, our analysis also reveals that diversity declined more sharply afterward (8–3 Myr) (Fig. 3), a time frame that overlaps with the arrival and diversification of placental carnivores. This rapid decline is likely the reason why our time series analyses estimated a significant effect of placental carnivores on the SA clade. Ultimately, however, our ability to find a definitive signal of competition in deep time is limited by the fact that there is rarely direct evidence for it in the fossil record (but see ^75^).

### Decaying negative impact of immigrant lineages

The impact of immigrant diversity on native taxa is more pronounced immediately following their arrival and during their initial diversification (Tables S3, S5, S7; Figs. S2-S7), while it decreases over time consistently across all our tests. This suggests a decrease in the intensity of competitive interactions, as native species possibly undergo selective extinction and immigrant descendants continue to evolve in the new environment and native species adapt, ultimately reaching a new equilibrium^13,16^. This decaying effect over time might explain the coexistence of SA native herbivores and immigrant herbivores from North America during the Late Pleistocene-mid Holocene (e.g., at least 7 and 4 sampled species of notoungulates and litopterns, respectively, were recorded), when both went extinct during the Megafauna extinction^76,77^.

Newly established immigrant lineages have the potential to affect the selective regime of native taxa, disrupting the diversification dynamics of resident clades^8,78^. However, the initial impact of newcomers could be most strongly expressed during the early stages of establishment and assimilation, before adaptation stabilizes the invader’s role^76^. Over the evolutionary timescale, the strength of interspecific competition can be diminished through spatial segregation, character displacement, and niche specialization^21,22^. If species specialize differently in resource use, interspecific competition weakens, reducing extinction rates and favoring the coexistence of multiple species^23^. Thus, although the short-term consequences of biological invasion are generally disruptive to the recipient fauna, interfering with established patterns of interaction among native species and potentially inducing their extinction in the long term, the net outcome over macroevolutionary time scales could be positive, increasing the diversity of the region^16,24^ by generating evolutionary radiations and ecological diversification or resource partitioning^12,46^.

### Andean uplift and diversity dynamics of South American mammalian faunas

The uplift of the Andes played a central role in shaping the diversity trajectories of native SA mammal groups (Fig. 4; Tables S4, S6, S8). Although this mountain-building process was diachronous and initiated in the Late Cretaceous, it intensified during the Neogene^80,81^, bringing profound changes to the continent’s topography and climate. These changes created new environments (e.g., montane forests, páramo, puna), reshaped existing habitats, and contributed to the expansion of open-arid ecosystems in southern South America^82,83^. Additionally, they facilitated the development of the Pebas system and the formation of the modern Amazon basin^82,84,85^.

Such environmental transformations are known to influence species turnover by driving both speciation through adaptation and ecological opportunity and extinction when species fail to cope with the new conditions^86^. Examples in South America include the radiation of sigmodontine rodents^87^, the evolution of hypsodonty in notoungulates^88^, and the contraction of platyrrhine monkeys’ latitudinal range in response to Andean uplift and climate cooling^86^. In contrast, some SA native groups, such as sparassodonts and notoungulates, experienced pronounced diversity declines during this period^36,41,74,60^.

Interestingly, the period of the intensified orogenesis during the Neogene^80,81^ coincided with significant fluctuations (i.e., declines and recoveries) in the species diversity trajectories of SA native groups, as well as immigrant groups from Africa, including platyrrhine monkeys and caviomorph rodents (Figs. 3, 4). These patterns reinforce the notion that Andean orogeny acted as a major driver of evolutionary change in South America by altering ecological landscapes and influencing the diversification and extinction dynamics of mammalian clades^24,61^.

### Alternative explanations and limitations

Antagonistic biotic interactions different from competition, such as predation and parasitism, may generate the same observed double-wedge pattern, dampening the diversity trajectory of natives as immigrant diversity increases^22,45^. Diseases carried by organisms may provide a strong mechanism for the competitive success of invaders^15,76^. While it may not be feasible to assess the impact of parasitism through the fossil record, we found no evidence for predator-naïve prey hypothesis (i.e., the arrival of placental carnivorans from North America) to explain the diversity decline trajectory of SA native herbivore lineages (i.e., litopterns and notoungulates combined species richness) (Fig. 4, Table S6) (but see ^60^ for an alternative explanation for toxodontids), suggesting that direct trophic interactions may not have been the primary driver in all cases.

Although our analyses suggest that competition may have driven biotic replacement, other explanations must also be considered. The correlated demise of native groups and rise of immigrants could reflect a differential response to physical factors, like environmental changes (e.g., climate shifts, habitat fragmentation)^42,83^. Although we found no association between native diversity trajectories and changes in the southern SA landscape, from dense vegetation to more open habitats, this abiotic variable is limited to a subregion of the continent, and its expression as a time series limits its ability to encapsulate the spatio-temporal nature. Hence, it does not reflect the evolution of habitats at a continental scale and their potential influence on SA mammalian diversity dynamics, as proposed by Vrba^42^. Furthermore, since the scope of our study was to analyze the responses of native fauna to biotic invasions, we did not examine the impact of physical factors on the diversity dynamics of immigrant groups. Consequently, passive displacement due to ecological vacancy—where natives decline independently and immigrants occupy emptied niches—cannot be ruled out entirely and may explain, in part, the observed patterns^37,38^. Large-scale evolutionary patterns are shaped by the interplay of physical and biotic processes ^4^. They may prevail or interact at different times and scales, making it challenging to disentangle. Ultimately, organisms and species interact with their environment and with each other; thus, both biotic and physical factors can act in concert to explain evolutionary change over time^62^.

The SA mammalian fossil record is characterized by its uneven spatio-temporal distribution^24,73,63^. Spatio-temporal biases in this record challenge the study of continent-wide diversity patterns and their underlying processes^43,64^. Additionally, different subregions of the continent may exhibit unique diversity patterns, making it challenging to determine the overall continental diversity dynamics^40,61^. In this study, we addressed this challenge using a novel method based on stochastic simulations and a deep learning model, which are robust to temporal and spatial heterogeneities in the fossil record^43,65^, allowing us to infer the diversity trajectories of the SA mammalian fauna throughout the Cenozoic at a continental scale. However, uncertainties in stratigraphic ranges, taxonomic assignments, and lineage-level ecology may remain. Moreover, our diversity proxies are based on species richness and do not account for functional, ecological, or phylogenetic diversity, which may reveal more nuanced interaction patterns.

Other challenges of our study include the fact that biotic interactions, such as competition, occur at local and regional scales, influencing population dynamics and geographic range limits over ecological time^66,67^. However, if the consequences of these interactions are consistent over time and across species’ geographic range, their effects may emerge at the species level and leave signatures at a macroevolutionary scale^54,68^. Increases in competition intensity decrease population sizes and intrinsic growth rates, thereby making species more vulnerable to stochastic extinction^6,66^. Moreover, the addition of new species may create new directed competition and intensify diffuse competition, leading to exclusion and increasing extinction probabilities^21,66,100^. The impact of competition on species distributions relies on coexistence^66^, besides the temporal coexistence of immigrants and native lineages on the SA continent (Figs. 3, 4), and despite the unavoidable incompleteness of the fossil record, they also co-occurred spatially; fossil occurrences of these taxa (i.e., placental and native predators; as well as native and immigrants herbivores) have been found in the same fossil locality and Formation (e.g., Tinguiririca, Salla, Cerro Azul, Chapadmalal, Tarija)^72,101–103^.

### Future directions and conclusion

While previous studies have identified diversity-dependent effects between distantly related clades with overlapping ecologies using hierarchical Bayesian models^8,10,104–106^, the possibility that these competitive effects diminish over time remains largely unexplored and should be considered in future model development. Expanding this framework to other continents and taxonomic groups would help assess whether the weakening influence of competition is a general macroevolutionary pattern. Additionally, incorporating trait-based approaches could clarify whether certain ecological traits (e.g., body size, dietary breadth, locomotor habits) made species more vulnerable to extinction or competitive replacement during biotic interchanges. Integrating paleoclimatic data with spatially explicit models would further refine our understanding of the relative roles of abiotic and biotic drivers^107^. Finally, genomic data from extant relatives could shed light on historical demographic responses to faunal turnover events, offering a more comprehensive view of long-term evolutionary dynamics.

In conclusion, using a quantitative framework, we showed that the arrival of mammalian lineages from other continents during the Cenozoic fundamentally reshaped South American faunas. While these immigrant clades contributed to increased overall diversity, they also played a role in the long-term decline of several native groups. The evolutionary history of South American mammals is thus marked by recurrent turnover events, driven by biotic interactions between native and immigrant taxa, rapid radiations of incoming lineages, and large-scale environmental transformations associated with Andean uplift. Together, our findings support the hypothesis that competition-driven biotic replacement is a key, yet temporally modulated, macroevolutionary force contributing to the dynamics of diversification and extinction shaping faunal assembly at continental scales.

## Methods

### Fossil occurrences data

We compiled a dataset of fossil occurrences for Cenozoic South American (SA) mammals at the species level (see Table S1 for a summary). This dataset includes the native SA groups: metatherians (Microbiotheria, Polydolopimorphia, Didelphimorphia, Paucituberculata, Sparassodonta, and “Ameridelphia”), xenarthrans (Pilosa and Cingulata), and native SA ungulates (SANUs: Notoungulata, Litopterna, Xenungulata, Pyrotheria, Astrapotheria, and Condylarthra), along with immigrant groups that arrived from Africa (Caviomorpha and Primates) and North America (Carnivora, Cricetidae, Artiodactyla, Perissodactyla, and Proboscidea). The dataset was compiled from the Palaeobiology Database (PBDB, paleobiodb.org, downloaded on August 20, 2023)—to which we contributed with ∼ 500 new entries (occurrences and taxonomic opinions)— the New and Old Worlds database of fossil mammals (nowdatabase.org, downloaded on June 6, 2023), and published literature^36,41,88,108^. Occurrences identified above the species level and non-terrestrial species were excluded. Records of the dromomerycine artiodactyls, peccaries, tapirs, and gomphotheres from the Late Miocene (∼6 Ma) in Amazonia^106,110^ were also excluded since the age of the records is controversial^111^. The compiled dataset was cleaned using a pipeline to identify potential errors related to space, time span, and taxonomy (including synonyms, misspellings, and duplicates) with the CoordinateCleaner and fossilbrush R packages^112,113^. The flagged occurrences were then inspected manually. The occurrence ages associated with the South American Land Mammals Ages (SALMAs) and some collections were updated with more recent dates^32,71,72,114–117^. We used the ‘look_up’ function and supplied the interval key argument with our updated ages implemented in palaeoverse^118^—occurrences with a temporal range larger than 15 million years were excluded^10^. The final dataset contains 6,214 occurrences spanning the last 66 million years, comprising 1,736 species across 673 genera sampled in the fossil record (Table S1, Dataset S1; see Supplementary Information).

### Diversity trajectories through time from fossil data

We used our compiled dataset of fossil occurrences to estimate the diversity trajectories of native and immigrant mammalian groups throughout the Cenozoic using the DeepDive method^43^. DeepDive is a software that employs stochastic simulations to generate diversity trajectories in space and time, which are then degraded to mimic the fossil record by applying preservation and sampling biases. The simulated data are then used to train a deep learning model based on long short-term memory (LSTM) neural networks, which learns to reconstruct the original biodiversity curves from the degraded fossil simulations. The trained models are then used to estimate biodiversity through time from the empirical fossil record, accounting for the temporal, spatial, and taxonomic heterogeneities observed in fossil preservation and sampling^43,65^. We utilized the autotune function implemented in DeepDive to ensure that the parameterization of the simulations accurately reflects the distribution of empirical data^65^ (Fig. S1, Table S1, Dataset S2; see Supplementary Information) and simulated 60,000 datasets for each analysis as training sets while using an additional 1,000 simulations as a validation set. We used 68 time bins, based on the boundaries of Cenozoic geological stages. Each stage was then further divided into smaller bins averaging 1 Myr in length, spanning from the beginning of the Danian (66 Ma) to the present. To account for spatial biases in fossil sampling, we assigned each occurrence to the following bioregions: tropical lowlands, extra-tropical lowlands, Pampas-Chaco, and central Andes (Fig. S1, Table S1). DeepDive’s simulations were therefore calibrated to reflect the sampling differences among these geographic units. Additionally, we conditioned the analyses on the modern diversity for the groups with extant diversity (Table S1). To account for dating uncertainty, we randomly resampled fossil ages 100 times and trained four models with different architectures (Table S2). The randomization and the predictions from the different models were combined to infer diversity over time and establish confidence intervals (CI) for each group.

### Correlation between diversity trajectories

We employed time series correlation analyses to test our hypothesis of negative biotic interactions, where the increasing diversity of immigrant groups drives a decline in native diversity, leading to biotic turnover and replacement (Fig. 1b, c). In addition, as this turnover pattern might be explained by immigrant and native lineages’ differential responses to environmental changes, we included two abiotic time series as additional predictors in our models: the Andes paleoelevation and the evolution of open habitats in southern SA^80,88,116^. The uplift of the Andes has been hypothesized to be connected to changes in the diversity dynamics of both extinct and extant SA mammals^36,74,87,60,61^, as its orogenic evolution has influenced regional climate and landscapes by affecting atmospheric circulation patterns, precipitation, and temperature regimes, and altering wind directions and river basins^84,120^. Whereas changes in the southern SA landscape, from dense vegetation to more open habitats, are considered to have driven the evolution of SA mammals, particularly herbivores^101,121^ (but see^88,122^).

Following the hypothetical framework outlined in Fig. 1, we tested the total diversity-dependence hypothesis considering: 1) a model with the combined diversities of immigrant groups from Africa and North America as predictors of total SA native diversity (i.e., combined species richness of metatherians, xenarthrans, and SANUs). Moreover, as competition is expected to occur within adaptive zones at the same trophic level^35^ we considered herbivores and carnivores groups separately modelling the following correlations: 2) To test the competition among herbivores and the predator-naïve prey hypotheses, we considered immigrant herbivore diversities from Africa (Caviomorpha) and North America (Artiodactyla, Perissodactyla, and Proboscidea) and immigrant predators (i.e., Carnivora order) from North America as predictors of the combined diversity of SA native herbivores (Notoungulata and Litopterna). 3) To test the competition among carnivores, we employed Carnivora and potential prey clades (SANUs, metatherians, Xenarthra, and Caviomorpha) diversities as predictors of SA native predators’ diversity (i.e., Sparassodonta).

We modeled generalized least squares (GLS) regressions with a first-order autoregressive process to account for temporal autocorrelation in the time series^123–125^, implemented in the nlme R package^126^. We generated a set of models (i.e., all possible combinations of biotic and abiotic time series predictors) for the three correlations tested (see above). Additionally, we modified the regression model by including an extra parameter, which we name the *decay effect*, that can capture a scenario where the impact of immigrant diversity is more pronounced at the time of their arrival, subsequently diminishing through time as their descendants assimilate in the new ecosystem and native species adapt, ultimately reaching a new equilibrium^13,16^. The *decay effect* parameter applies an exponential distribution to transform the diversity trajectories of immigrants, with the initial values in the transformed time series (i.e., at the time of dispersal of the immigrant clade) having more weight, whereas subsequent values have exponentially smaller weights. Specifically, the transformed trajectories at time *t* are a function of the ratio between the diversity trajectory at time *t*, i.e., *v_t,_* and the diversity trajectory at time *t* – 1:

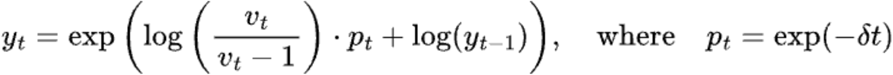

where δ ≥ 0 is the decay parameter. When δ = 0, the shape of the transformed curve is identical to the original trajectory, while δ > 0 leads to early fluctuations (i.e., when *t* is small) being stronger, while later fluctuations are flattened out. The resulting vector of values *y_t_* was z-transformed using the scale function in R prior to running all regressions, and across all δ values. We optimized the value of the *decay effect* as a free parameter in our model, through a grid search in a maximum likelihood framework (Figs. S3, S5, S7). We then used bias-corrected Akaike’s information criterion (AICc) and AICc weights to quantify model fit (Tables S3, S5, S7). We used model averaging to account for uncertainties in model selection^127,128^ and computed model-averaged regression coefficients for the top-ranking models with cumulative AICc weights = 0.65. We obtained the upper and lower bounds of the 65% confidence intervals for each parameter using the R package MuMIn^126^ (Dataset S3; see Supplementary Information).

### Data availability

All datasets and code analyses will be available after the peer review process.

## Acknowledgments

We want to thank all paleontologists whose hard work collecting and describing fossil specimens has generated and published South American fossil mammal data used in this study, as well as those who have contributed to incorporating these data into the Paleobiology Database.

